# Antagonistic mechanisms of probiotic *Aliivibrio* sp. strain Vl2 against *Moritella viscosa*: Evidence from co-cultivation and transcriptomic analysis

**DOI:** 10.1101/2024.12.21.629903

**Authors:** Marius Steen Dobloug, Stanislav Iakhno, Simen Foyn Nørstebø, Henning Sørum

**Affiliations:** Department of Paraclinical Sciences, Faculty of Veterinary Medicine, Norwegian University of Life Sciences, Elizabeth Stephansens vei 15, 1430 Ås, Norway; Previwo AS, Ullevålsveien 68, 0454 Oslo, Norway

## Abstract

Winter ulcers are a significant challenge to the Norwegian aquaculture industry and the primary causative agent is *Moritella viscosa*. Control measures are limited due to the lack of effective vaccines and limited use of antibiotics, reflecting the global effort to combat antibiotic resistance. It was recently demonstrated that probiotic *Aliivibrio* spp. colonize the skin and ulcers of Atlantic salmon and their association with a reduced prevalence of winter ulcers. These observations suggest that *M. viscosa* and *Aliivibrio* spp. may interact within ulcers *in vivo*. In this study, we have investigated how the probiotic *Aliivibrio* sp. strain Vl2 modulates *M. viscosa in vitro*, using both co-cultures and CHSE cell cultures. We found that the probiotic strain exhibits antagonistic effects against *M. viscosa*, reducing its growth and pathogenicity toward salmonid cells. Transcriptome analysis of *Aliivibrio* Vl2 revealed potential mechanisms impeding the growth of the competing pathogen. Together, our findings demonstrate how the probiotic bacterium can inhibit *M. viscosa in vitro* and suggests potential mechanisms that may explain the reduced prevalence of winter ulcers observed in the field.

## 1. Introduction

Bacterial ulcers are a major challenge for Norwegian open pen aquaculture of Atlantic salmon (*Salmo salar*) and the prevalence has increased substantially the last decade^1^. This is primarily due to winter ulcers, a disease occurring most frequently during the winter months, typically causing round ulcers on the lateral sides of the fish. In 2023, outbreaks were documented in 320 different Norwegian farms^1^ and the actual prevalence is likely even higher, as it is relatively easy to diagnose macroscopically and not a notifiable disease.

*Moritella viscosa* is the main aetiological agent causing winter-ulcer disease^2^*. M. viscosa* is more adherent when cultured at 4 °C compared to 15 °C which has been proposed as a reason for its increased pathogenicity in winter, when water temperatures are lower^3,4^. The pathogen does not invade host cells, as demonstrated by immunohistochemical staining of infected tissues and cell-culture infection models^3,5^. Tunsjø *et al*. proposed that cell damage is caused by extracellular proteins secreted by the bacteria, which leads to cytoskeleton disruption, pore formation, and ultimately cell lysis^3^.

*M. viscosa* is often isolated from winter ulcers together with other bacteria^6,7^. One of these agents, *Aliivibrio wodanis,* has been shown to strongly reduce the growth and acute virulence of *M. viscosa* ^8^. However, *A. wodanis* is a pathogen by itself, and a co-infection generally leads to a prolonged course of illness^9^.

Despite bacterial infections constituting a major challenge in the Norwegian salmon farming industry, there is an overall low use of antibiotics^1^. This is partly due to effective vaccines, along with strict regulatory oversight and biosecurity measures^10,11^. Vaccines for *M. viscosa*, however, have failed to adequately protect the Atlantic salmon from winter ulcers, leaving the salmon farming industry vulnerable. This situation opens the door for alternative health management options such as probiotic bacteria, bacteriocins, bacteriophages and other novel interventions^12,13^.

The relationship between vertebrates and resident bacteria dates back hundreds of millions of years and serve a wide variety of functions in the host^14^. In fish, microbial communities have been studied in skin, gills and gut with several links made to disease resistance^15^. Beneficial bacteria introduced to the host are often referred to as probiotics, defined as *“live microorganisms which when administered in adequate amounts confer a health benefit on the host”* ^16,17^. Probiotic bacteria have been shown to serve many different purposes in fish, such as control of infectious diseases^18–20^, water quality improvement^21,22^ and functional ingredients^23,24^. These benefits can be conferred through competitive exclusion of pathogens^25–28^, quorum quenching^29^, immunomodulation^30^ and more^31,32^. Previously, studies have documented the associations between probiotic *Aliivibrio* spp. and reduced prevalence of winter ulcers in Atlantic salmon^33^. We have recently recovered these probiotic species from skin and co-cultures in ulcerated tissues following bath administration (manuscript in preparation). The mechanisms by which these probiotics exert their effects remain unknown.

To characterize these mechanisms, we first conducted co-cultivation experiments to study interactions between *M. viscosa* and *Aliivibrio* sp. strain Vl2. We then evaluated the cytotoxic potential of the *Aliivibrio* strain toward a CHSE cell line and examined whether its culture supernatant could mitigate the harmful effects of *M. viscosa*-on these cells. Finally, we characterized the transcriptomic profile of *Aliivibrio* Vl2 during this interaction using RNA sequencing (RNA-seq).

## 2. Materials and methods

### 2.1 Strain collection

Three different isolates of *M. viscosa* (strain 7 (from a strain collection isolated at Stokkasjøen in 2019), NVI-3632 (RefSeq genome accession no. GCF_900120025.1) and NVI-5427 (RefSeq genome accession no. GCF_900120285.1)), all originally isolated from ulcerated fish were collected from the freeze stock (−80 °C) at the Norwegian University of Life Sciences (NMBU) Department of Paraclinical Sciences, Bacteriology and Mycology Unit. These isolates represent the three clonal complexes (CC) of *M. viscosa* that has been isolated from farmed fish in Norway, where CC1 and CC3 are currently reported to dominate^34^. Strain NVI-3632 is the classic strain belonging to the CC1 category, while strain NVI-5427 belongs to CC2 and strain 7 belongs to the CC3 category.

*Aliivibrio* sp. strain Vl2, NCIMB 42592 was originally isolated, cultivated and stored as detailed by Klakegg *et al*.^33^ and collected from freeze stock (−80 °C) at Previwo AS (Oslo, Norway).

All bacteria were cultured as following unless otherwise specified: Bacteria were taken from stock at −80 °C and first cultured on blood agar plates (OXOID, CM0271, blood agar base No. 2, with 5 % bovine blood and 0.9 % NaCl) for five days at 10 °C. A starter culture was prepared by transferring single colonies from a blood agar plate to Luria–Bertani (LB) broth (Merck, Germany) with 0.9 % NaCl added. The cultures were incubated at 8 °C with 120 rpm for three days before expansion 1:10, followed by incubation for three more days.

### 2.2 Co-cultures in LB broth

Starter cultures of *Aliivibrio* Vl2*, M. viscosa* (strain 7, NVI-5427 and NVI-3632) were prepared as described above. In two different experiments, growth of *M. viscosa* and *Aliivibrio* Vl2 was assessed in liquid co-cultures and compared to monocultures. Each liquid co-culture experiment was conducted in duplicate.

First, we investigated if the growth of *M. viscosa* (strain 7) and *A*.Vl2 was altered in liquid co-cultures at equal concentrations, compared to their growth in monocultures. Ten ml of *M. viscosa* (strain 7) at 0.5 OD_600_ was added to 50 ml Erlenmeyer vials together with 10 ml of *Aliivibrio* Vl2 at 0.5 OD_600_ (co-culture). The same volumes of the same bacterial cultures were also added to 10 ml of 0.9 % LB broth without bacteria (monoculture). All cultures were incubated at 8 °C and 120 rpm for five days, with daily samplings for serial dilution and plating to determine colony-forming units (CFU) per millilitre (ml) of liquid culture.

In the second experiment, we assessed the inhibitory effects of *Aliivibrio* Vl2 on all three strains of *M. viscosa*. To show the probiotic’s inhibitory activity, the temperature and the relative proportion of probiotics in the co-culture were reduced. To set up the co-cultures, 10 ml of *M. viscosa* culture (OD_600_ = 0.5) and 10 ml of Vl2 culture (OD_600_ = 0.05) were combined in 50 ml Erlenmeyer vials.

Monocultures were prepared using the same concentrations of each bacterial culture, added to 10 ml of 0.9 % LB broth without other bacteria. The respective cultures were kept at 120 rpm at 4 °C for five days and sampled daily for serial dilution and plating. The growth was evaluated based on the number of colony-forming units (CFU) per ml of liquid culture.

### 2.3 Co-cultures on blood agar plates

*M. viscosa* (strain 7) and *Aliivibrio* Vl2 were cultured as previously described. Both strains were then crossed with each other on a fresh blood agar plate by streaking one strain horizontally and the other vertically, creating an intersection between the two bacteria (shown in Figure 2). The plate was incubated at 10 °C for six days, after which the interactions between the two species were evaluated.

Next, a starter culture of *Aliivibrio* Vl2 was prepared as previously described before transfer to 50 ml Falcon tubes. To isolate supernatants, the tubes were centrifuged at 4600x g for 10 minutes and the supernatants were double sterile filtered with 0.2 µm pore size Minisart Syringe Filter (Sartorius, Germany), before storage in aliquots at −80 °C until use.

*M. viscosa* (strain 7) was transferred from a blood agar plate and uniformly spread onto fresh blood agar plates using a Retro C80 plate carousel (Montebello diagnostics, Norway). Twenty microliters (µl) of *Aliivibrio* Vl2 supernatant thawed from −80 °C storage was spotted onto the spread bacteria as a single droplet. The plates were then incubated at 4 °C for seven days before evaluation.

### 2.4 Scanning electron microscopy (SEM)

Five days old colonies of *Aliivibrio* Vl2 were transferred to 10 ml fixative consisting of 5 ml 4 % paraformaldehyde, 2.5 ml 0.5 M PIPES, 0.5 ml 25 % glutaraldehyde and 2 ml dH_2_O. Fixated bacteria were incubated on a glass slide coated with 1 mg/ml poly-L-lysin, dehydrated in an alcohol gradient, critical point dried and sputter-coated with platinum particles. Bacteria on prepared slides were inspected with a Zeiss EVO 50 scanning electron microscope (20 000x magnification, 10 kV EHT, 10-15 pA probe). The same was carried out for bacterial extracellular vesicles (BEVs), prepared and isolated as described below (See Cell culture infection assay).

### 2.5 Spatially separated co-cultures

For spatially separated co-cultures, semi-permeable 25 mm diameter regenerated cellulose bags with MW cutoff 12-14 kDa (Spectra/Por, Los Angeles, CA) were prepared after the method of Colquhoun *et al.* ^35^. Three knots were used to seal the bottom end of each bag and a sterile Venofix IV-tube (with cap and without needle, B. Braun, Germany) was inserted. The top of the bag was sealed and secured with knots using a separate sterile IV-tube and the sealed bags were disinfected using 70 % ethanol.

Five ml 0.5 OD_600_ *M. viscosa* (NVI-3632) or *Aliivibrio* Vl2 was added to respective bags through the IV-tube (Figure 1). In one bag of *M. viscosa* (NVI-3632), 50 µl of *Aliivibrio* Vl2 BEVs was also added (See Cell culture infection assay for preparation and isolation). Each bag was submerged in 100 ml 0.9 % NaCl LB broth in 500 ml Erlenmeyer vials. In addition, one vial contained both a bag of *M. viscosa* NVI-3632 and *Aliivibrio* Vl2. In total, this yielded four cultures: *M. viscosa* monoculture, *Aliivibrio* Vl2 monoculture, Vl2 and *M. viscosa* co-culture, and *M. viscosa* with *Aliivibrio* Vl2 BEVs. Each bag was incubated at 8 °C with 120 rpm and sampled for OD_600_ measurement at day 0, 1, 2, 3, 7 and 11. The LB broth outside the bags in each vial was inspected visually and cultured on blood agar plates to assess contamination at each timepoint. The experiment was performed in duplicate.

**Figure 1:**
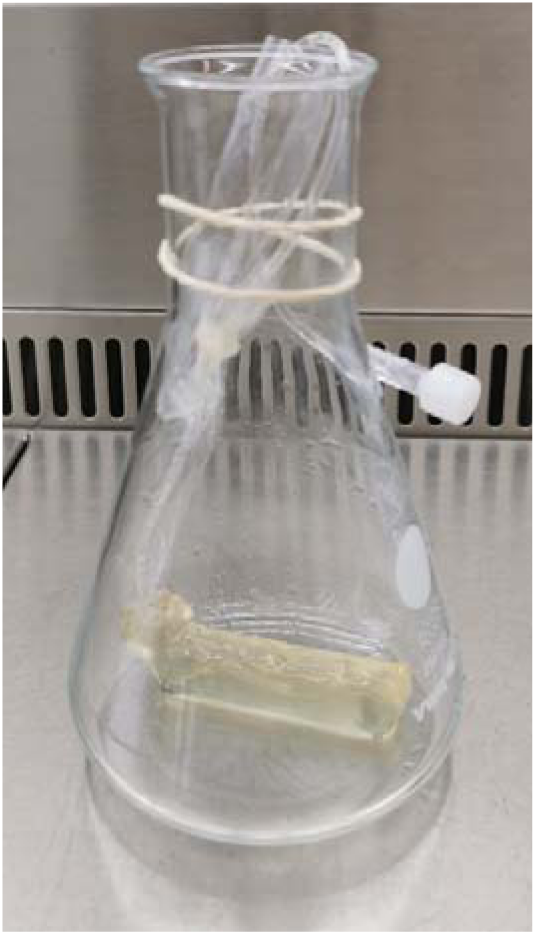
Sealed autoclaved semipermeable bag with sterile IV-tube inserted containing 5 ml of 0.5 OD_600_ *M. viscosa*in a 500 ml Erlenmeyer vial, prior to addition of 100 ml 0.9 % NaCl LB broth.

**Figure 2:**
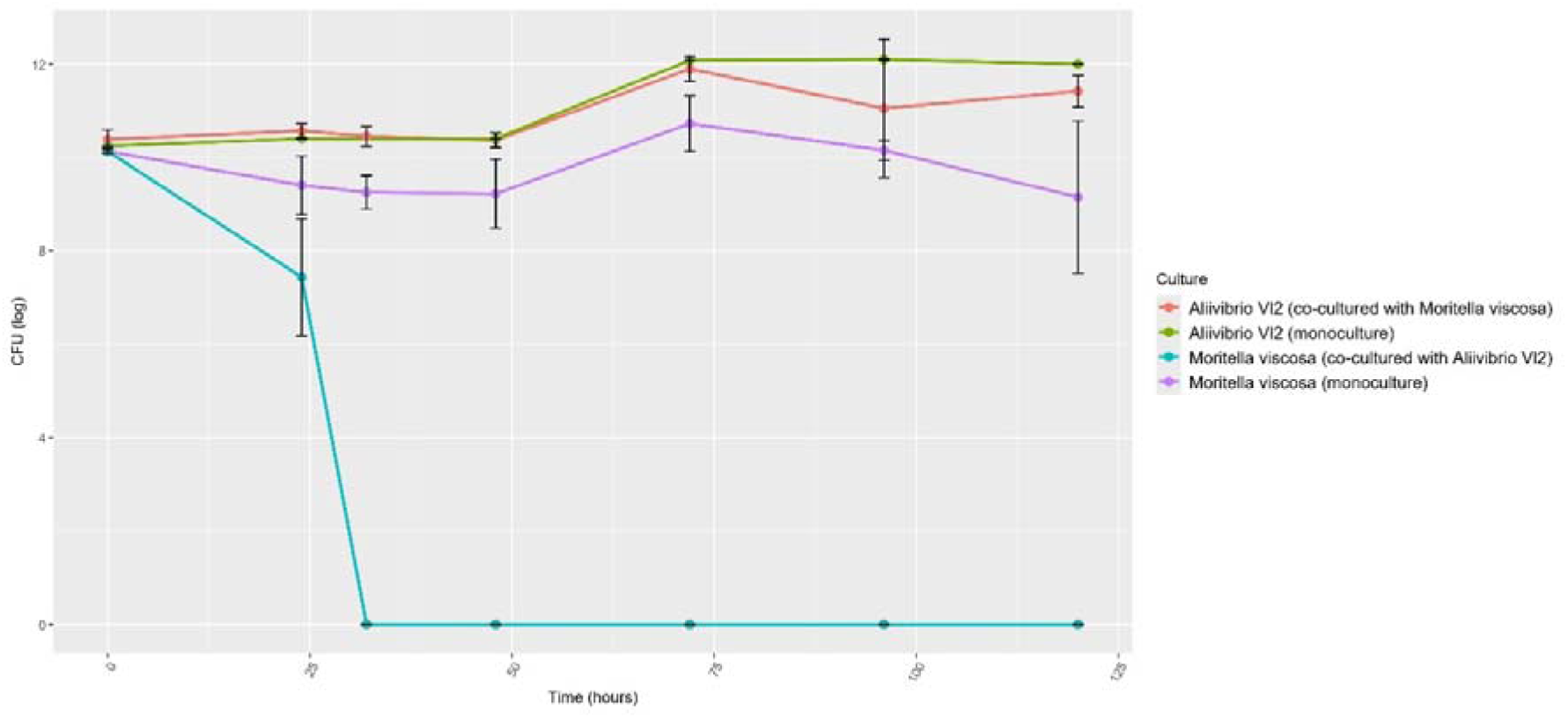
Mean CFU/ml (log) with standard deviations (SD) of *M. viscosa*(strain 7) and *Aliivibrio*Vl2 when co-cultured in equal concentrations at 8 °C (n=8). Figure created in Rstudio.

### 2.6 Cytotoxicity assay

The supernatant from *Aliivibrio* Vl2 and *M. viscosa* NVI-3632 was harvested as previously described for supernatant growth inhibition, but not sterile filtered, resulting in a mixed bacterial culture containing supernatant and approximately 10^3^ cfu/ml. The cultures were then aliquoted and stored at −80 °C until use.

CHSE-214 salmon cell line (Merck, Germany) was grown in 75 cm^2^ culture flasks with LB-15 + glutamax (Gibco, USA) and 10 % FBS media (Gibco, USA) at 20 °C. Cells were detached with 0.05 % trypsin (Gibco, USA) and split weekly at ≈80 % confluence in a 1:4 ratio. Cells were counted with a TC20 Automated Cell Counter (Bio-Rad Laboratories, USA) and viability was assessed with erythrosine B (Logos biosystems, South Korea). One hundred microliters of fresh media containing approximately 20 000 cells at passage 20-30 with >97 % viability was transferred to each well in 96-well plates and incubated at 20 °C. When the wells reached ≈80 % confluence, the cytotoxicity assays were conducted by adding 20 µl of the mixed bacterial culture to duplicate wells and incubated at 8°C for 24 hours.

To evaluate toxic effects on the cells, all wells were inspected visually using an EVOS M5000 Imaging System (Thermofisher, USA) and measured for LDH concentration using the CyQUANT™ LDH cytotoxicity assay kit (Thermofisher, USA). Wells were then carefully washed five times with 1x PBS (Gibco, USA) and incubated with fresh media containing 200 µg/ml gentamycin for four hours, before measuring cell viability with Alamar Blue (Thermofisher, USA) as per the manufacturer’s instructions using a Cytation 3 Cell Imaging Multi-Mode Reader (BioTek, USA). Wells without CHSE cells were used to subtract background signal. Two biological replicates were used per group in this experiment.

### 2.7 Cell culture infection assays

CHSE cells were prepared as described above. Bacterial cultures of *M. viscosa* strain 7 and *Aliivibrio* Vl2 were prepared as described above and adjusted an OD_600_ of 0.5. Ten ml of each of these cultures were combined and co-cultured for three days. The culture was then expanded 1:10 and incubated for an additional three days, followed by a second 1:3 expansion and three more days of incubation, before harvest of the supernatant (Supernatant 1). A monoculture of *Aliivibrio* Vl2 without *M. viscosa* was also prepared in the same manner (Supernatant 2). Both supernatants were harvested by centrifugation at 15 000x g at 4 °C for 10 minutes, double sterile filtered with 0.2 µm pore size Minisart Syringe Filter, and aliquoted to Eppendorf tubes. In addition, bacterial extracellular vesicles (BEVs) were isolated from 300 ml Supernatant 1 by ultracentrifugation as described by Brudal *et al.* ^36^. The BEVs were suspended in 600 µl PBS and aliquoted into eppendorf tubes. This procedure was repeated for another 300 ml of supernatant 1, but the BEVs were suspended in 6 ml PBS, introducing a 1:10 dilution. All supernatants and BEVs were stored at −80 °C until further use. To verify the absence of culturable bacteria, the supernatant and BEV samples were cultured on 0.9 % NaCl blood agar plates. Presence of BEVs was also controlled by SEM as described in the section Scanning electron microscopy.

The cell culture infection assay was conducted on CHSE cells with ≈80 % confluency by adding 20 µl of *M. viscosa* (strain 7) at 0.8(±0.1) OD_600_ and 20 µl of either PBS, *Aliivibrio* Vl2 supernatant or *Aliivibrio* Vl2 BEVs. For negative controls, 40 µl PBS were added to the CHSE cells. After incubation at 8 °C for 18 hours, each well was carefully washed five times with PBS and incubated with fresh media containing 200 µg/ml gentamycin for four hours, before measuring cell viability with Alamar Blue (Thermofisher) as per the manufacturer’s instructions. Each group was replicated in four wells (technical replicates) per plate, and the experiment was repeated four times (biological replicates). Each plate contained four wells per group. Wells exceeding the detection limit were excluded, resulting in the omission of one well. Cell viability was calculated by subtracting the background signal from wells without CHSE cells, and the mean value for each group was compared to the controls. Visual assessment of cell damage was performed using the EVOS M5000 Imaging System (Thermofisher, USA), with images captured from a fixed location using the Cytation 3 Cell Imaging Multi-Mode Reader (BioTek, USA).

### 2.8 RNA-seq analysis

Monocultures and co-cultures of *Aliivibrio* Vl2 and *M. viscosa* (NVI-3632) in LB broth (0.9 % NaCl) were prepared as described for liquid co-cultures. Four replicate co-cultures consisting of 10 ml *Aliivibrio* Vl2 at 0.5 OD_600_ and 10 ml *M. viscosa* (NVI-3632) at 0.5 OD_600_ were set up, in addition to four monocultures of *Aliivibrio* Vl2 at 0.5 OD_600_ in 10 ml 0.9 % NaCl LB broth. All eight cultures were incubated at 8 °C and 120 rpm for 72 hours, with samples collected at 0, 24, 48 and 72 hours for serial dilution and determination of CFU/ml (Table S1 and Figure S1). The same experiment was then repeated for preparation of samples for RNA sequencing, with all cultures sampled after 24 hours.

The cultures were centrifugated at 4600x g, and the supernatant was discarded. The bacterial pellets were resuspended and RNA was extracted using the RNeasy Protect Bacteria Mini Kit (Qiagen, Germany) following the manufacturer’s instructions. RNA concentration and purity was measured using a Multiskan Sky Spectrophotometer (Thermofisher, USA), and RNA integrity (RIN) was evaluated with a 4200 TapeStation (Agilent, USA). Samples were sent to Novogene (UK) for library preparation, including rRNA removal and prokaryotic mRNA paired-end sequencing with Illumina PE150, generating 5-10 million reads per sample for biological quadruplicates of both groups.

The Raw sequencing reads were quality filtered using “bbduk”, version 37.48 (BBMap – Bushnell B., https://sourceforge.net/projects/bbmap/) and Filtlong v0.2.0 (https://github.com/rrwick/Filtlong), respectively. Hybrid (short and long reads) whole-genome assemblies were obtained with Unicycler v0.4.8^37^. The quality of the assembled genomes was assessed by mapping the sequencing reads to the *de novo* assembled genomes using Minimap2 and samtools ver.1.3.1 as well as by the assessment of the assembly graphs. The assembled genomes were annotated using Prokka^38^.

The transcriptome reference file was obtained using an AGAT conversion Perl script, “agat_convert_sp_gff2gtf.pl”, to obtain the GTF file from the Prokka-generated GFF file (REF: Dainat J. AGAT: Another Gff Analysis Toolkit to handle annotations in any GTF/GFF format. (Version v0.7.0). Zenodo. https://www.doi.org/10.5281/zenodo.3552717). The reads were aligned using STAR to the annotated genome using the genome sequence (from the de novo assembly) and the GTF file as input^39^.

The obtained gene counts were processed using the DESeq2 package^40^ in RStudio (Version 2024.04.2+764). Normalization was performed using DESeq2’s built-in median ratio method, which accounts for sequencing depth and library size differences, and normalized counts were extracted. For visualization, log2 transformation and variance-stabilizing transformation (VST) were applied to stabilize variance across gene expression levels. The normalized and transformed data were visualized using boxplots to ensure consistency across samples. The DESeq2 pipeline was subsequently applied to prepare for differential expression analysis, including size factor estimation and dispersion modeling.

The OmicsBox bioinformatics suite (formerly Blast2Go^41^) was used for the Gene set enrichment analysis (GSEA) filtered for 0.05 P-value and 1000 permutations. The sequence data is available at Fairdomhub (DOI: https://doi.org/10.15490/fairdomhub.1.datafile.7605.1)

### 2.9 Statistics

The cytotoxicity assay was analysed by a nonparametric Mann-Whitney test by comparing the cell viability per well for each group. Damage reduction was analysed by unpaired two-tailed t-test by comparing the average cell viability per plate for each group. All assumptions have been fulfilled. All data reported are untransformed values.

## 3. Results

### 3.1 Co-cultures in LB broth

When co-cultured with *Aliivibrio* Vl2 in equal concentrations, the number of detected *M. viscosa* (strain 7) colonies was reduced by 2 logs after 24 hours compared to the *M. viscosa* monoculture (Figure 2). No colonies of *M. viscosa* were detected after 32 hours. The growth of *Aliivibrio* Vl2 was less affected by *M. viscosa,* but the number of colonies was reduced by 1 log after 96 hours and 0.6 log after 120 hours, when compared to the *Aliivibrio* Vl2 monoculture (n=8).

When evaluating the growth of three strains of *M. viscosa* co-cultured with *Aliivibrio* Vl2 in a 10:1 ratio, all strains were outcompeted by Vl2 within 120 hours (Figure 3). Notably, no colonies from *M. viscosa* strain 7 were detected after 72 hours. Strain NVI-3632 was strongly reduced after 72 hours and not detectable after 120 hours. Strain NVI-5427 appeared less impacted by the probiotic species during earlier sampling points but was undetectable at the final sampling. *Aliivibrio* Vl2 appeared to be less affected by the presence of any *M. viscosa* strain as the growth in monoculture was similar to the co-cultures.

**Figure 3:**
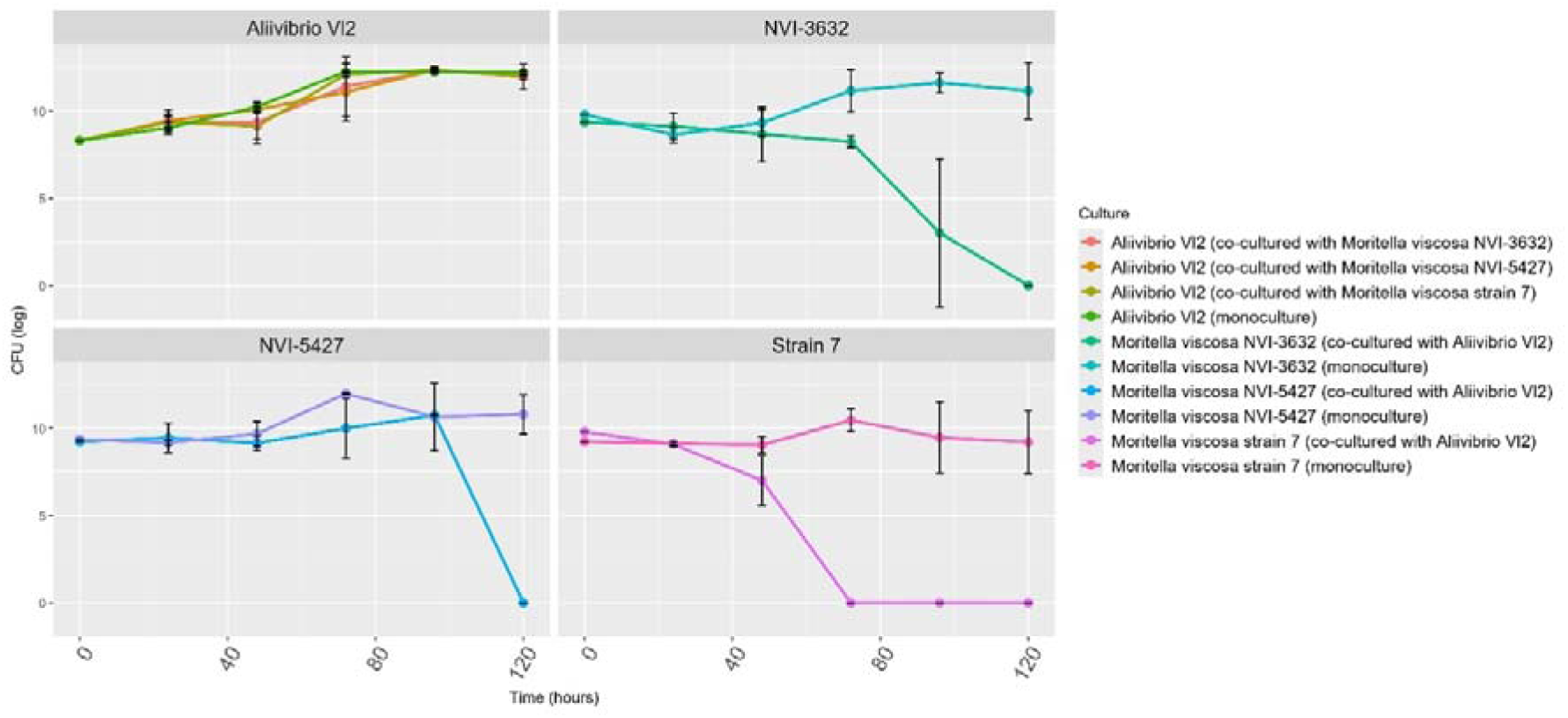
Mean CFU/ml (log) with SD of three different *M. viscosa*strains and Aliivibrio Vl2 cultivated in a 10:1 ratio as co-cultures and as monocultures at 4 °C (n=20). Figure created in Rstudio.

### 3.2 Co-cultures on blood agar plates

After six days, the *Aliivibrio* Vl2 exhibited growth extending over *M. viscosa* on both sides (Figure 4A). When exposing *M. viscosa* to supernatant from *Aliivibrio* Vl2, reduced growth was observed after seven days in the area where the supernatant had been placed (Figure 4B).

**Figure 4:**
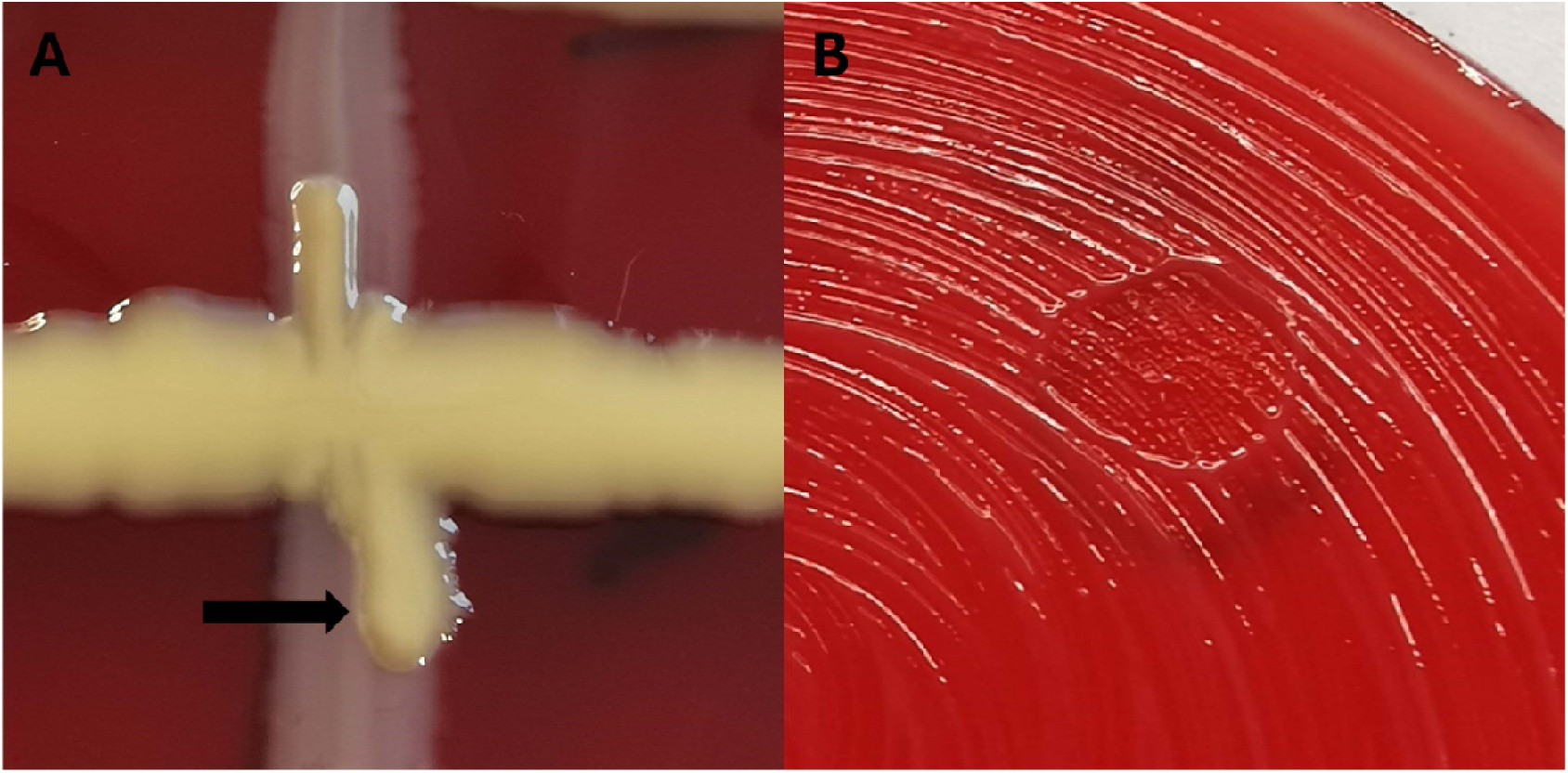
Interactions on blood agar plates. A) Cross streak of Aliivibrio Vl2 (horizontal streak) with zone lines (arrow) in the contact area against M. viscosa (vertical streak). B) Growth reduction of M. viscosa by Aliivibrio Vl2 supernatant. Figure created in Rstudio.

### 3.3 Scanning electron microscopy

Scanning electron images visualize vegetative Aliivibrio Vl2 with surrounding vesicles (Figure S2A) and vesicles isolated by ultracentrifugation from Aliivibrio Vl2 (Figure S2B).

### 3.4 Spatially separated co-cultures

Spatially separated co-cultures of M. viscosa and Aliivibrio Vl2 showed clear differences in growth between co-cultures and monocultures for both bacteria (Figure 5). The growth of M. viscosa was lower at every sampling point past day 0 when co-cultured with Aliivibrio Vl2 compared to when monocultured. By day 11, there was an average reduction in growth of 53.1 %. There was also a reduction of 19 % in Aliivibrio Vl2 growth when co-cultured with M. viscosa as compared to a monoculture. There was no apparent growth reduction of M. viscosa when exposed to Aliivibrio Vl2 vesicles.

**Figure 5:**
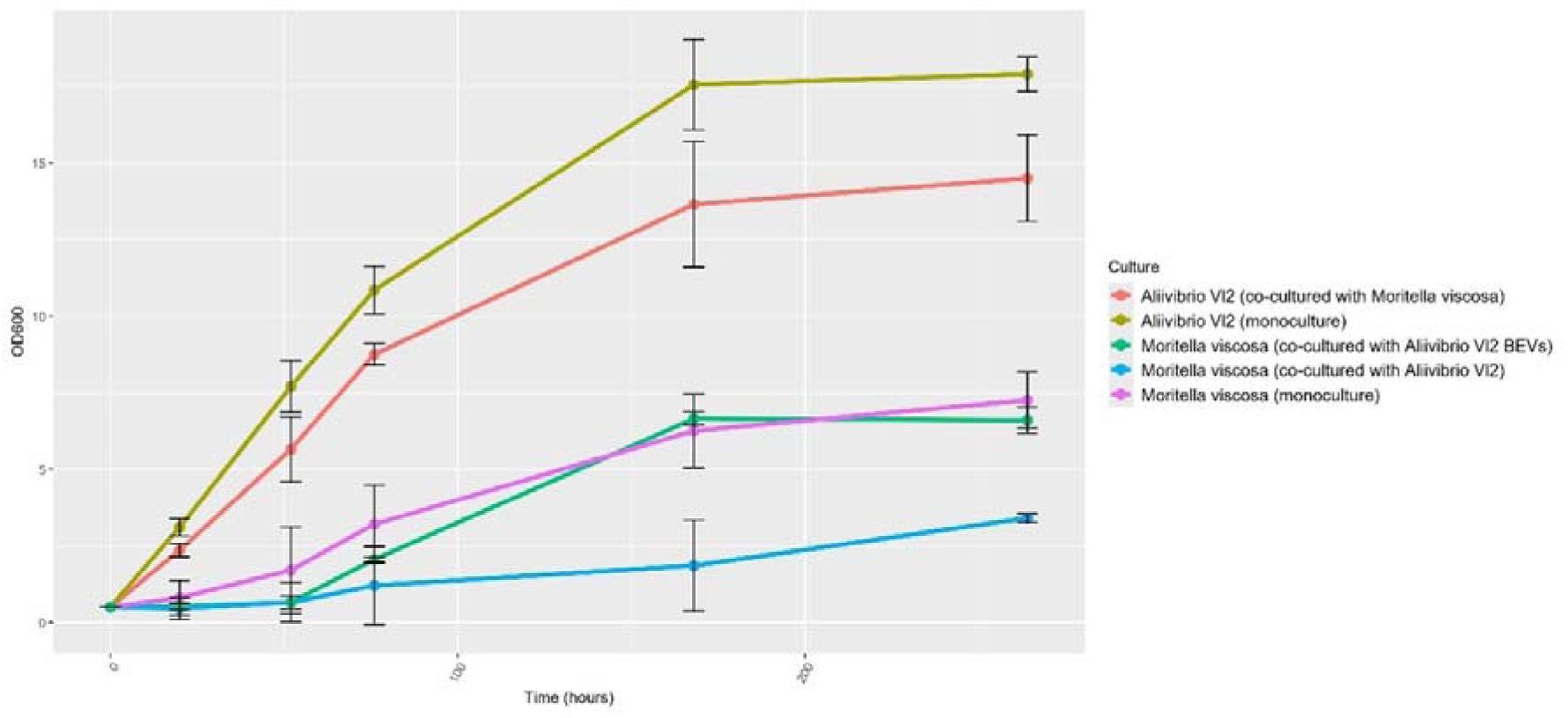
Mean OD_600_ with SD of *M. viscosa*(NVI-3632) and *Aliivibrio*Vl2 cultivated in semipermeable bags, either as monocultures in separate vials or as co-cultures in shared vials. Duplicate bags also contained *M. viscosa*(NVI-3632) with *Aliivibrio*Vl2 BEVs (n=10). Figure created in Rstudio.

### 3.5 Probiotic cytotoxicity assay

*Aliivibrio* Vl2 showed no visible signs of damaging CHSE cells, and cell viability, as measured by Alamar blue was comparable to the control (Figure 6) (P>0.05 by nonparametric Mann-Whitney test). In contrast, *M. viscosa* reduced viability of the CHSE cell culture to 46 % in 24 hours, a significant difference compared to the control group (P=0.022 by nonparametric Mann-Whitney test).

**Figure 6:**
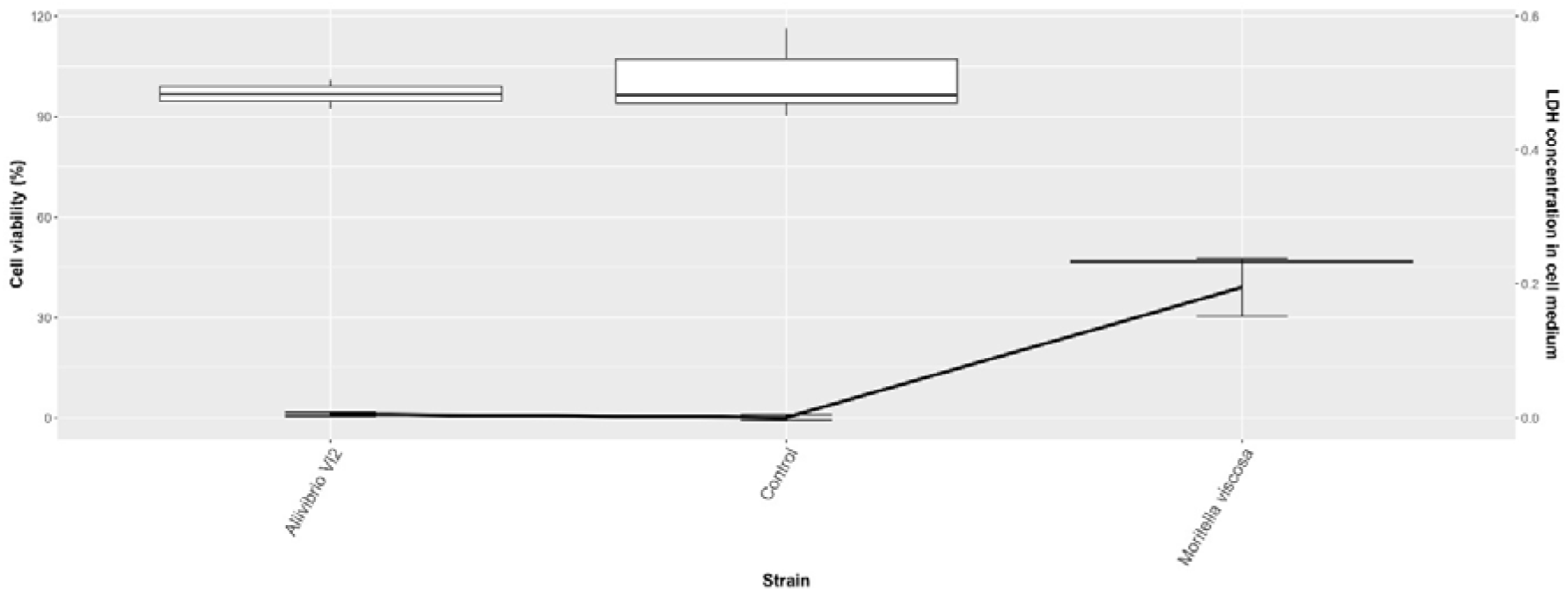
Cell viability (box plot) and LDH concentration (line plot) measured in vitro after 24-hour incubation of CHSE cell cultures with *Aliivibrio*Vl2, *M. viscosa*(strain 7) or control. The boxplots represent the median values along with interquartile ranges, while the line plot represents the mean values with SD (n=6). Figure created in Rstudio.

Additionally, LDH release measured by relative fluorescent units (RFU), was substantially higher in wells with *M. viscosa* than in those exposed to *Aliivibrio* Vl2 or in the controls.

### 3.6 Cell culture damage reduction assay

Cell viability in wells containing only *M. viscosa* was significantly lower than in wells with *M. viscosa* and *Aliivibrio* Vl2 supernatant 1 (from co-culture, P<0.0001) or supernatant 2 (from monoculture, P=0.002) (Figure 7). Visually, more live cells were observed in wells with containing probiotic supernatant compared to those with only *M. viscosa* (Figure 8). However, signs of cell damage typically seen prior to cell death were present in wells with supernatant, such as elongation and rounding. Addition of *Aliivibrio* Vl2 BEVs did not significantly reduce the damage caused by *M. viscosa* as measured by cell viability (P>0.05), and no damage reduction was observed visually. The wells with added supernatant 1 had a higher level of cell viability in average compared to supernatant 2, but they were not significantly different from each other (P>0.05).

**Figure 7:**
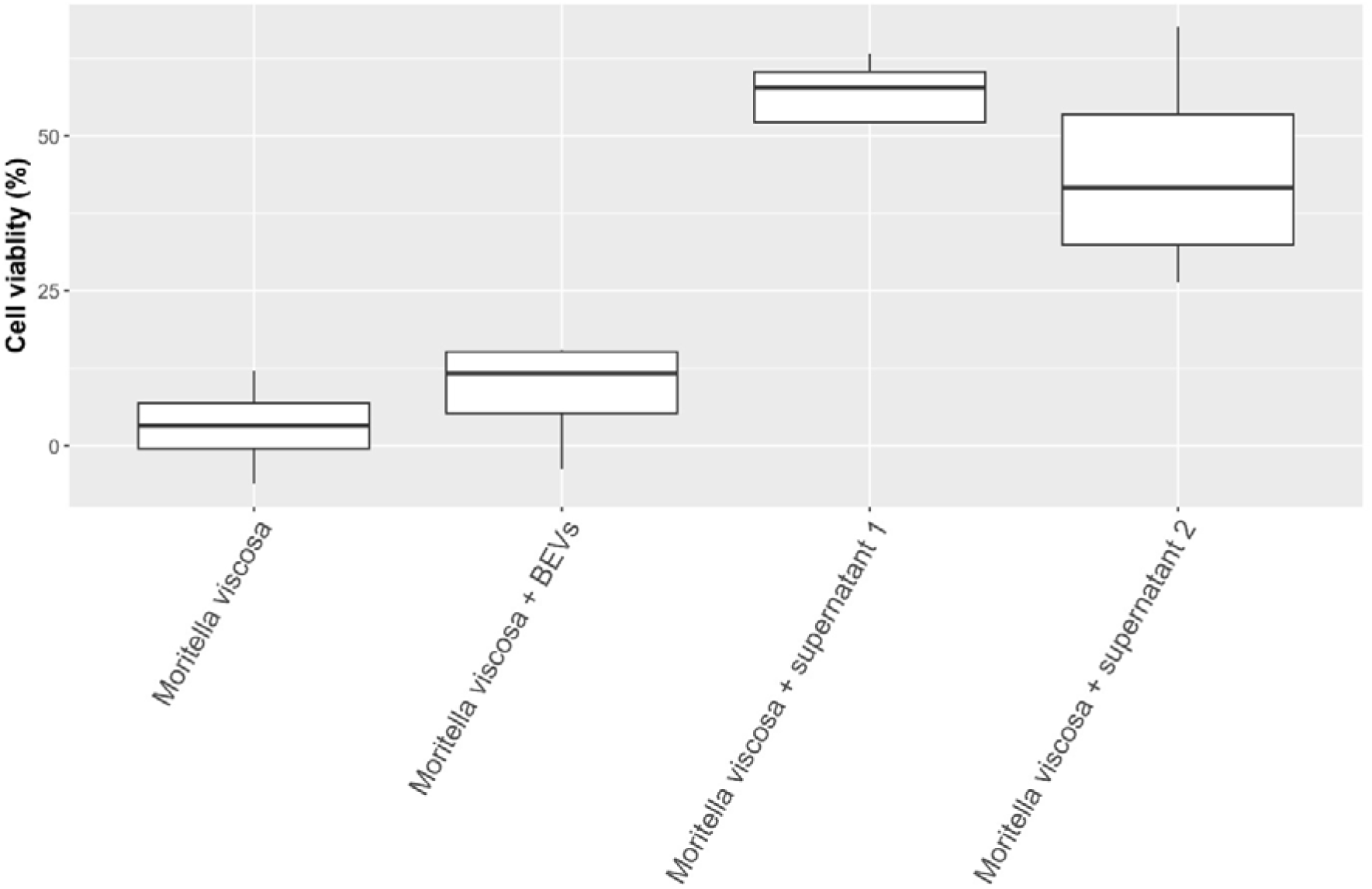
Cell viability (% reduction relative to control) after exposure to M. viscosa (strain 7) and probiotic supernatant or BEVs from Aliivibrio Vl2. The boxplots represent the median values along with interquartile ranges (n=20). Figure created in Rstudio.

**Figure 8:**
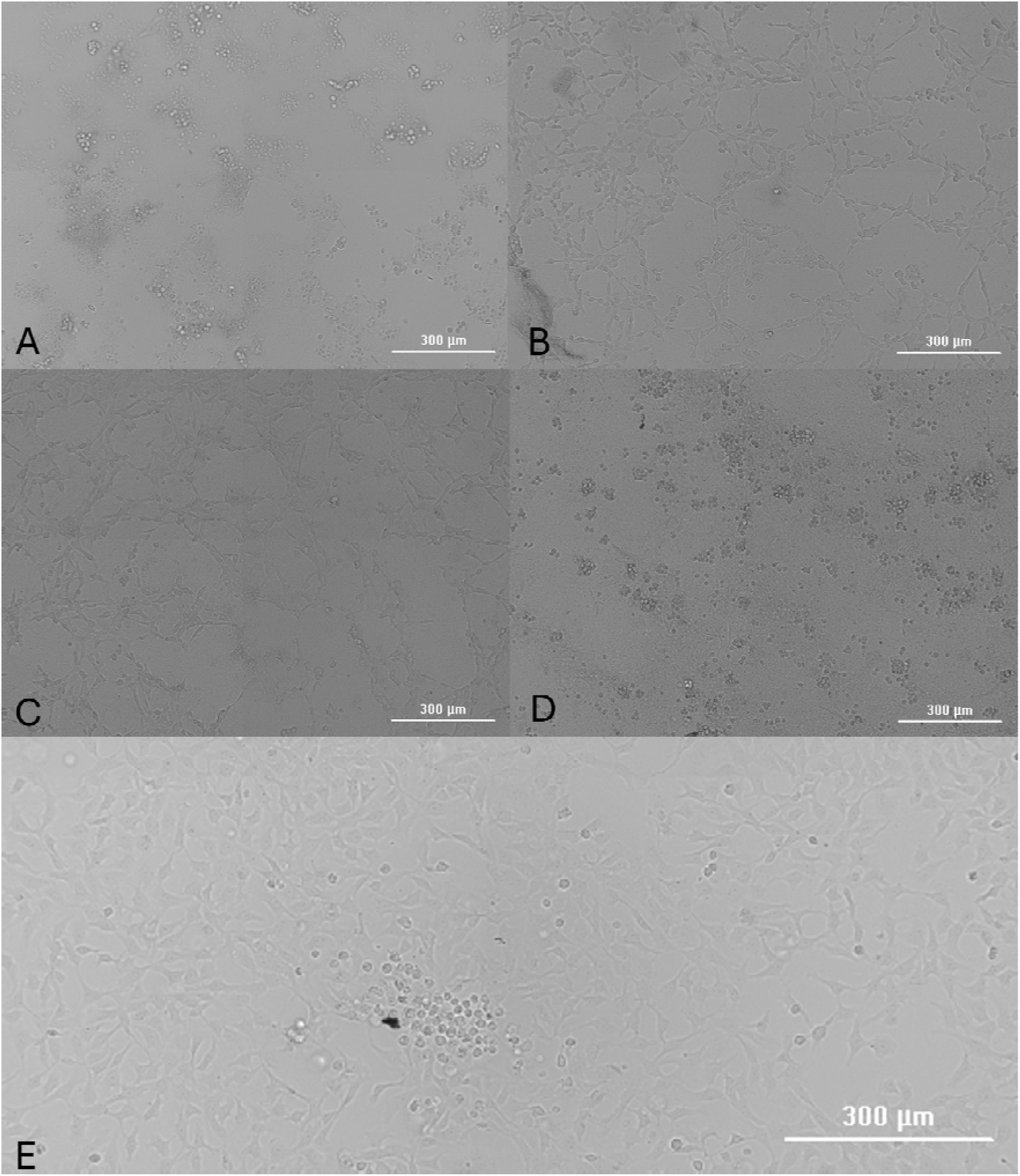
Representative images of CHSE cells after exposure to (A) M. viscosa only, (B) M. viscosa and Aliivibrio Vl2 supernatant 1, (C) M. viscosa and Aliivibrio Vl2 supernatant 2, (D) M. viscosa and Aliivibrio Vl2 BEV’s and (E) control.

### 3.7 Transcriptomics

The growth recorded in monocultures and co-cultures is presented in Table S1 and Figure S1. A total of 140 355 970 transcripts annotated to 3 870 different genes were recognized for Aliivibrio Vl2 overall. Of these, 708 genes were significantly differentially expressed when comparing the Aliivibrio Vl2 monoculture to the co-culture with M. viscosa (P_adj_<0.05). Of these significant differentially expressed genes (DEGs), 326 were upregulated and 382 were downregulated in the co-culture. The 100 most significant DEGs were used to compare the samples, presented in Figure 9.

**Figure 9:**
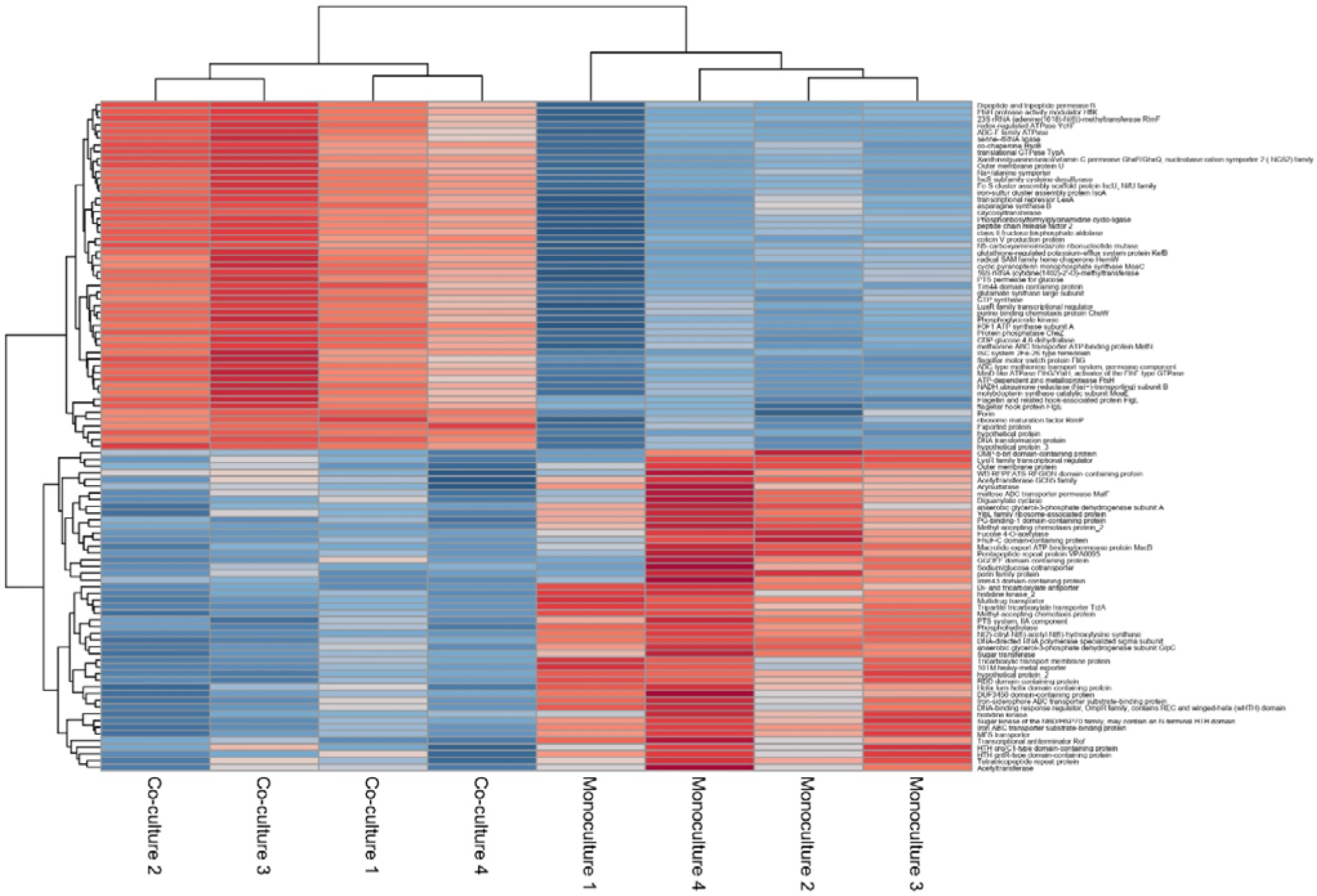
Heatmap with dendrograms of the 100 most significant DEG’s with co-cultures and monocultures in separate vertical clusters based on Euclidian distance (n=8). Figure created in Rstudio.

Using BLAST^42^, the DEGs were functionally annotated and assigned to 712 Gene Ontology (GO) terms. In the co-culture, downregulated GO terms (651) significantly outnumbered the upregulated (63) ones. The 32 most significantly upregulated and 32 most significantly downregulated GO terms are presented in Figure 10.

**Figure 10:**
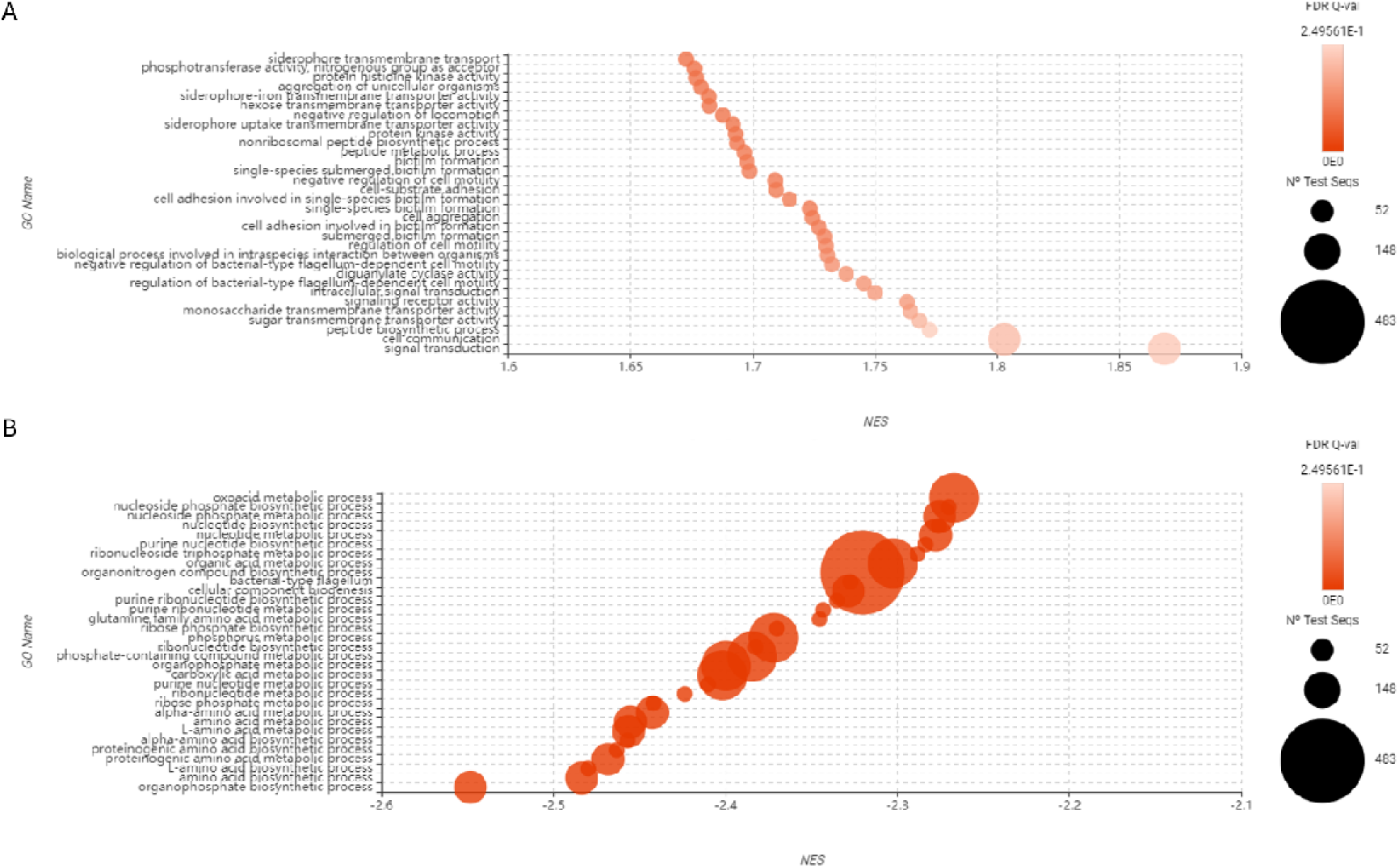
Functional enrichment analysis based on the sequences recognized by BLAST(n=8). GO names are sorted based on Normalized Enrichment Score (NES) in descension for up-regulated genes (A) and ascension for down-regulated genes (B). Figure created in Omicsbox.

Most downregulated GO terms were associated with metabolic and biosynthetic processes for nucleotides and amino acids, while the upregulated GO terms were more varied in their functional classifications (Figure 10). Several significantly upregulated genes were associated with production, secretion and uptake of siderophores (Table 1). This siderophore activity, in conjunction with upregulated cell signalling and biofilm formation, may modulate the probiotic’s competitive advantage over M. viscosa. Alterations in motility-related gene expression were also observed, with flagellar-genes downregulated and genes negatively regulating motility upregulated.

**Table 1:**
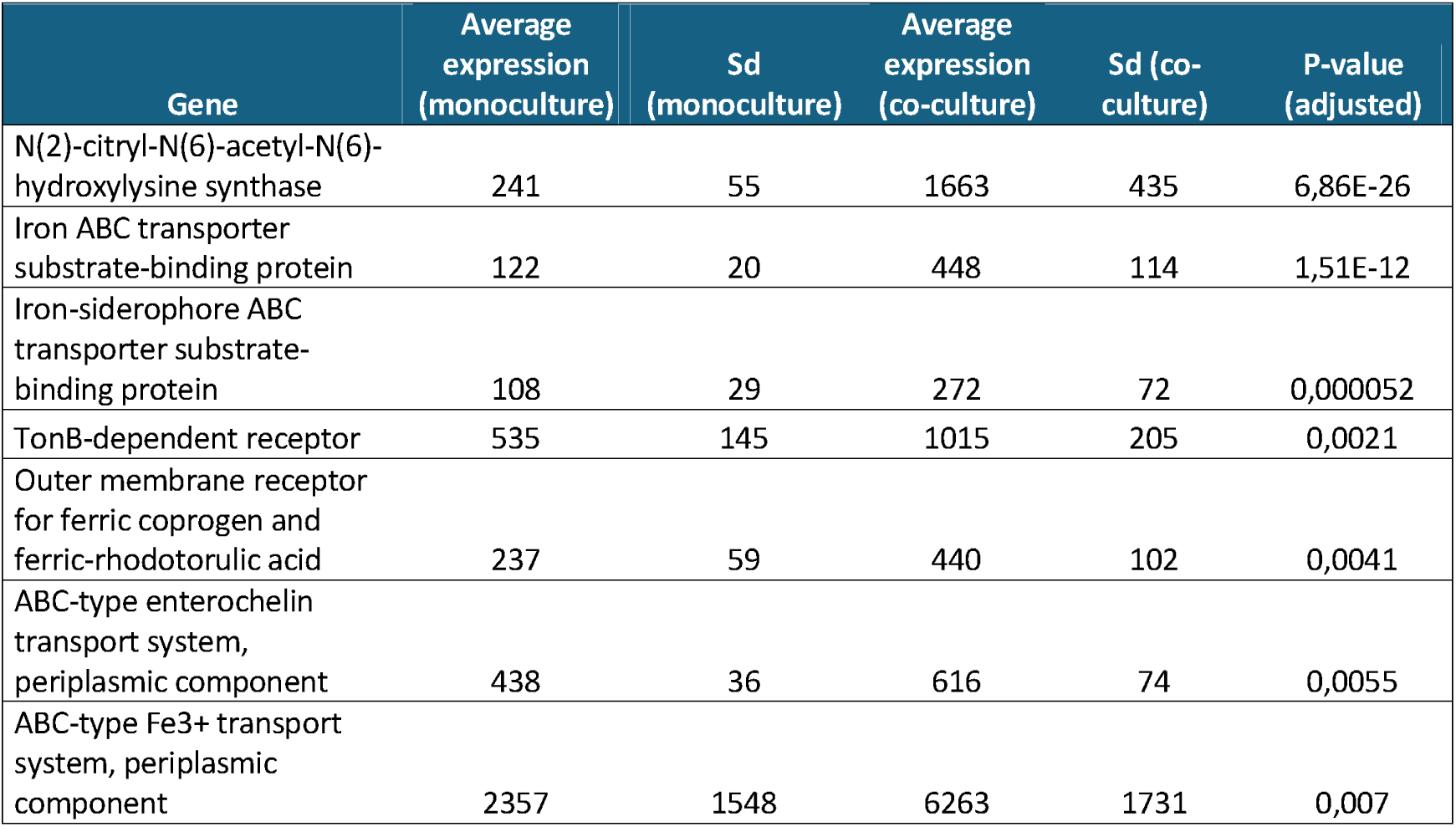
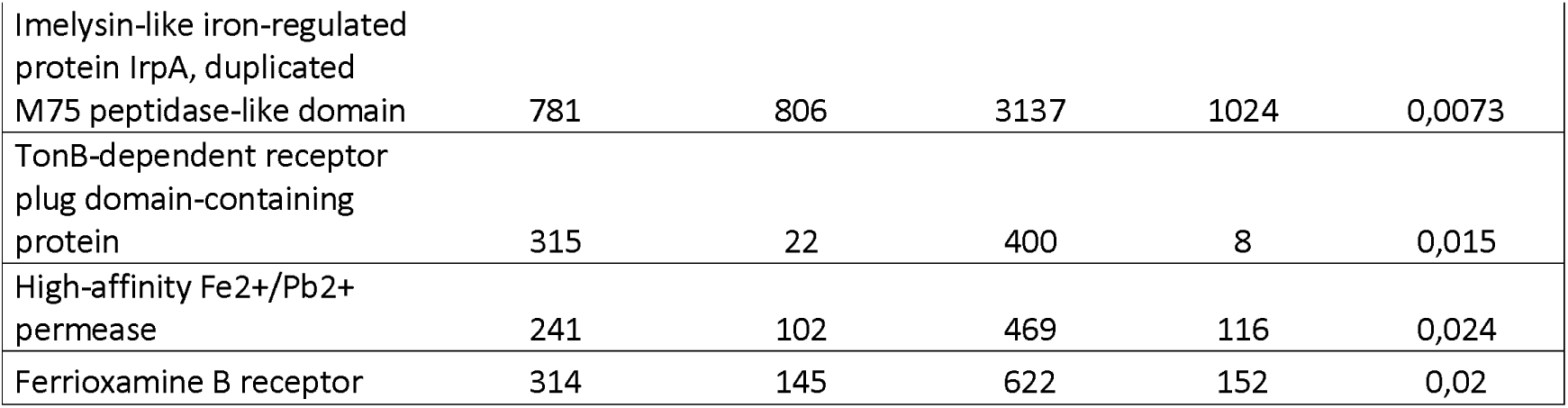
Upregulated genes (P_adj_<0.05) for siderophores.

## 4. Discussion

In the present study we have investigated how two naturally occurring bacteria in Atlantic salmon, the ulcer pathogen *M. viscosa* and the probiotic species *Aliivibrio* sp. strain Vl2, interact under different conditions *in vitro*. Our findings show that growth of *M. viscosa* is significantly inhibited by *Aliivibrio* Vl2, suggesting a competitive relationship driven by the probiotics mechanisms to outcompete the pathogen. These mechanisms, which appear to be contact-independent, also reduce the level of damage inflicted on CHSE cell cultures by *M. viscosa*.

### Co-cultures in liquid and solid media

When *M. viscosa* and *Aliivibrio* Vl2 were co-cultured in equal concentrations at 8 °C, *M. viscosa* was not detectable after 28 hours. When reducing the concentration of the probiotic strain by tenfold and lowering of the temperature to 4 °C, the growth of three different *M. viscosa* strains was still impeded when co-cultured with the probiotic *Aliivibrio* species. There are clear differences in how long the different strains of *M. viscosa* were able to resist the presence *Aliivibrio* Vl2, although the end-result was the same (Figure 3). The difference in susceptibility may be related to the different CC groups. However, as we only used one strain per CC group, these observations may be strain specific.

Bacteria can develop specific responses when encountering other bacteria on solid media. Plate crossing experiments have previously been carried out for *M. viscosa* and *A. wodanis*, with documented contact-independent inhibition of *M. viscosa*. Hjerde *et al*. attributed the inhibition to increased gene expression level within a bacteriocin locus and siderophore systems^8^. Here we observed a different phenomenon, with overgrowth of *Aliivibrio* Vl2 and contact-dependant inhibition of *M. viscosa*. Zonal lines at the contact point suggests that the probiotic bacteria produce distinct substances not found elsewhere in the colony^43^. This overgrowth further indicates that the probiotic bacterium is unaffected by the presence of *M. viscosa* and may even be attracted to it, possibly stimulated by substances produced by the pathogen - a phenomenon previously documented in other bacterial interactions^44,45^.

Finally, we investigated if the growth reduction was contact dependent. When the two bacteria were co-cultured within individual semi-permeable bags, the growth of *M. viscosa* was still strongly reduced (Figure 5). Hence, the inhibition is non-contact dependent, which is also supported by the growth inhibition seen on blood agar plates by the addition of probiotic supernatant (Figure 4B).

### Cell cultures

Mixed cultures of *Aliivibrio* Vl2 and *M. viscosa* were screened for cytotoxic effects (Figure 6). While the probiotic caused no damage to the CHSE cells, *M. viscosa* reduced the cell viability to 46 % compared to that of the controls. Additionally, the presence of *M. viscosa* elevated LDH level in the culture medium, indicating cell lysis, a response not observed with *Aliivibrio* Vl2.

When CHSE cells were exposed to *M. viscosa* in a damage reduction assay, the cell survival increased from 3.8 % to 49.4 % in average if probiotic supernatant 1 (from co-culture) or 2 (from monoculture) was also added to the wells (Figure 7). This indicates that *Aliivibrio* Vl2 can effectively reduce the damage to CHSE cells during *M. viscosa* infections through secretion of inhibitory compounds into its supernatant. Although the cell survival increased drastically by the presence of both probiotic supernatants, most cells were still negatively affected by the presence of *M. viscosa*, evident by widespread cell elongation and some cell rounding (Figure 8), which are signs of stress typically seen prior to detachment and cell death^33^. This suggests that the probiotic supernatants may mitigate the stress caused by *M. viscosa*, potentially prolonging cell survival despite the observed signs of cellular distress.

When comparing the effects of the two supernatants on the cell survival no apparent difference was detected. This means that the production of inhibitory compounds by *Aliivibrio* Vl2 was not dependent on the presence of *M. viscosa*. However, supernatant 1 resulted in more cells surviving in most experiments, implying that *Aliivibrio* Vl2 may have adjusted its extracellular production repertoire to specifically target *M. viscosa* during co-culture.

### BEVs

Many bacterial species continuously release vesicles into the surrounding environment, and some vesicles from probiotic bacteria have been shown to harbour anti-bacterial properties in fish^46^. While we observed the production of BEVs in *Aliivibrio* Vl2, these BEVs did not affect the viability of *M. viscosa* when *Aliivibrio* BEVs were added directly to the semi-permeable bags used for cultivation (Figure 5) or when introduced to cell cultures (Figure 7 and Figure 8).

### Transcriptomics

The transcriptome analysis of *Aliivibrio* Vl2 revealed a significant downregulation of genes associated with various metabolic and biosynthetic pathways, particularly those involved in protein synthesis. We propose that during the encounter between *Aliivibrio* Vl2 and *M. viscosa*, the former may reprioritize its core metabolic activities in favour of functions more relevant to adaptation and survival within a co-culture environment.

Motility-related processes appeared to be restricted in the co-cultured *Aliivibrio* Vl2, as genes associated with flagellar assembly were downregulated, while those involved in the negative regulation of motility showed increased expression. Instead of prioritizing motility-related gene expression, the co-cultured aliivibrio exhibited increased expression of genes associated with cell adhesion, aggregation, and biofilm formation. Additionally, the expression of several genes linked to siderophore activity was upregulated. These changes in the global gene expression suggest that *Aliivibrio* Vl2 may be initiating survival-associated pathways in response to the presence of *M. viscosa*.

The downregulated genes were associated with multiple GO terms with major overlaps between them (bubble sizes, Figure 10B). This is likely due to these transcripts being applicable to several similar metabolic processes, while the upregulated genes presented in Figure 10A appear to be associated with effects in which the specific transcripts are exclusively linked to individual pathways. When experiencing environmental stressors, bacteria can gather on a surface and encase themselves in protective biofilm^47–49^. The population must communicate through cell signals to coordinate the production of biofilm matrix^50^, which is also represented by several upregulated GO-terms in our data set (Figure 10A). Additionally, standalone DEGs, such as acyl-homoserine-lactone (AHL) synthase being central to quorum sensing, were found to be upregulated.

Iron is essential for the growth and survival of nearly all bacteria. However, due to its limited bioavailability, many bacteria produce siderophores to acquire iron. Siderophores are molecules with high affinity for iron, released by bacteria into the surrounding environment to scavenge iron and transport it back. Many different siderophore systems have been documented over the years^51^. In this study, we observed the inhibitory effects of *Aliivibrio* Vl2 on the viability of *M. viscosa* in co-cultures and documented significant upregulation of siderophore pathways in aliivibrio cells during this interaction. Although a direct causal link cannot be confirmed, these findings imply that *Aliivibrio* Vl2 competes for available iron by producing siderophores, thereby depriving *M. viscosa* of this essential growth factor. As siderophores are small (usually 500-1000 Da), they are able to pass through the semi-permeable tubings and have contact-independent antagonistic effects, as observed in these studies.

*Aliivibrio* species have previously been shown to produce siderophores to drive competitive exclusion in co-cultures with other species^52,53^. Eickhoff *et al*. showed that the siderophore known as aerobactin allows *Aliivibrio fischeri* to establish a niche by denying growth of its competitor.

Aerobactin is biosynthesized from lysine with three intermediary steps before aerobactin synthase finally generates the siderophore (KEGG pathway map00997). Aerobactin synthase was not significantly differentially expressed in our dataset, but the final intermediary enzyme, N(2)-citryl-N(6)-acetyl-N(6)-hydroxylysine synthase, was the most significantly differentially expressed gene in the entire transcriptomics dataset (P_adj_ = 6,86E-26) with a fold-change of 6.9. The co-culture was sampled daily for 72 hours, with inhibition of *M. viscosa* starting at 48 hours (Table S1 and Figure S1). To capture transcripts relevant to the observed interactions, the samples taken at 24 hours were selected for transcriptomic analysis. The subsequent strong inhibition of *M. viscosa* likely coincides with the completion of aerobactin biosynthesis with the secretion of substantial quantities into the environment. In addition to competitive exclusion via resource competition, aerobactin can enhance bacterial defences in biofilm formation and oxidative stress protection^54^.

Some bacteria can steal iron sequestered by siderophores produced by other bacteria, a competitive strategy known as siderophore piracy^55^. Even unculturable bacteria can be stimulated to grow when co-cultured with a siderophore-producing bacterium through piracy^44^. Although there is a spike in aerobactin-precursor transcripts, the final step is not (yet) upregulated. Despite this, several TonB-receptors and ABC-transporters important for siderophore-complex import, are significantly upregulated. Upregulated genes in four additional siderophore systems were recognised in the co-cultures. These were specific receptors responsible for uptake of ferric coprogen, rhodotulic acid, enterochelin and ferrioxamine. However, the genes related to production of these siderophores were not detected in *Aliivibrio* Vl2. Although this can be due to uncharacterized annotations, we speculate that the probiotic bacteria may be a siderophore pirate, capable of scavenging siderophore complexes produced by other species. Notably, *M. viscosa* is known to produce siderophores^56^ which could be pirated by *Aliivibrio* Vl2. This phenomenon may also explain the observed attraction towards *M. viscosa* on blood agar plates.

Competitive exclusion can also be exerted through secretion of antimicrobial substances^57,58^. There are several genes encoding the production of such molecules, which were expressed by *Aliivibrio* Vl2. Genes encoding antibiotic biosynthesis monooxygenase and Isopenicillin N synthase with related dioxygenases are involved in the production of potent antibiotics. Several genes linked to toxin and anti-toxin systems were also upregulated in the co-cultured *Aliivibrio* Vl2, indicating an activation of multiple offensive and defensive mechanisms that can benefit the probiotic in the competition with *M. viscosa*.

## Conclusion

In Norwegian salmon aquaculture, it is not uncommon for *M. viscosa* induced winter ulcers to be co-infected with *A. wodanis* ^9^. This pathogenic *Aliivibrio* species exhibits a robust non-contact-dependent inhibitory effect on *M. viscosa*, likely mediated by bacteriocins and siderophore production^8^. Recent work has shown that probiotic *Aliivibrio* spp. also colonize salmon ulcers, potentially reducing their prevalence through competitive exclusion (manuscript in preparation).

In this study, we have shown that *Aliivibrio* sp. strain Vl2 exhibits antagonistic traits against *M. viscosa* that resemble those of *A. wodanis* but with potential mechanisms that resemble those of *A. fischeri*. The observed effects are likely linked to the expression of genes involved in siderophore production, synthesis of antibacterial compounds, and biofilm production. These findings provide a foundation for further investigation into the mechanisms underlying these interactions, guided by the transcriptome analysis presented here.

## Supplementary materials

Table S1, CFU/ml in cultures over time; Figure S1: CFU/ml in cultures over time. Red arrow indicates timepoint of sampling for RNA-seq.

## Author contributions

Marius Steen Dobloug: Conceptualization, Data curation, Formal analysis, Investigation, Methodology, Visualization, Writing – original draft, Writing – review and editing. Stanislav Iakhno: Investigation, Methodology, Writing – review and editing. Simen Foyn Nørstebø: Methodology, Supervision, writing – review and editing. Henning Sørum: Conceptualization, Investigation, Methodology, Supervision, Writing – review and editing.

