## Supplementary materials for "Antagonistic mechanisms of probiotic *Aliivibrio* sp. strain Vl2 against *Moritella viscosa*: Evidence from co-cultivation and transcriptomic analysis"

Table S1: CFU/ml in cultures over time.

| **Culture** | **start** | **24t** | **48t** | **72t** |
| --- | --- | --- | --- | --- |
| M.Viscosa NVI-3632 | 11,9138139 | 12,2148438 | 14,2304489 | 12,845098 |
| Aliivibrio balderis | 12,4771213 | 12,60206 | 14,3579348 | 14,0253059 |
| Kokultur balderis | 12,4771213 | 12,544068 | 14,30103 | 14,3180633 |
| kokultur MV | 11,9138139 | 11,5185139 | 10 | 0 |

Figure S1: CFU (log)/ml in cultures over time. Red arrow indicates timepoint of sampling for RNA-seq.

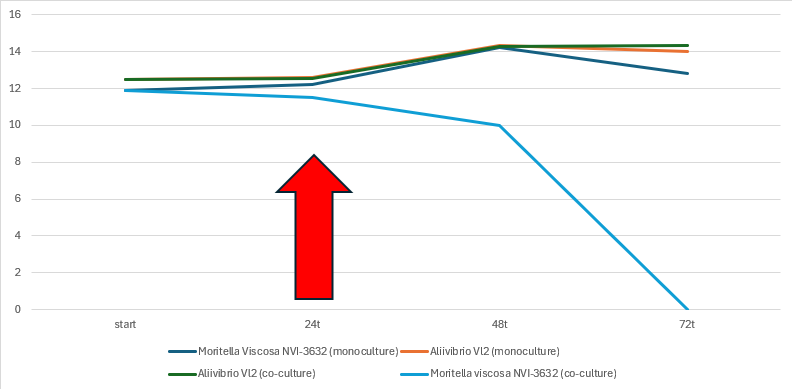

Figure S2: SEM imaging at 20 000 x magnification of Ca. *A.* *balderis* (A) colony and (B) isolated BEV’s.

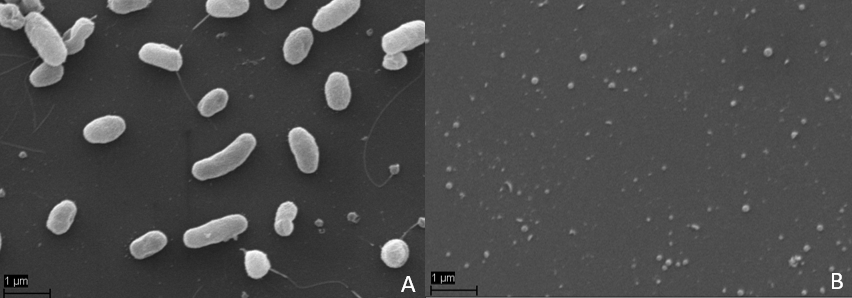
